# Gene mutations and molecular evolution of the chikungunya virus E1 and E2 from 2014 to 2019

**DOI:** 10.1101/2022.03.16.484685

**Authors:** Xiaoxia Li, Xinyue Li, Jinyue Liu, Song Liang, Hanzong Chen, Changwen Ke, Chengsong Wan

## Abstract

Chikungunya virus (CHIKV), a mosquito-borne *Alphavirus*, is the etiological agent of chikungunya fever. To investigate the prevalence and genetic characteristics of CHIKV in China, we performed a molecular epidemiological study during the 2014–2019 period. Phylogenetic analysis of CHIKV in the 2014-2019 shows that it is currently clustered geographically, which means that CHIKV’s local adaptability is gradually strengthening. Focusing on CHIKV adaptive mutation E1-A226V is not obvious in our study. The CHIKV E1 amino acid non-synonymous mutation results revealed the common mutation site M343I. Most mutation sites belong to the American epidemic strains, and the mutation rate is 2.8%. In 31 sequences, 19 non-synonymous mutations were found in CHIKV E1 (total mutation rate 20/499 = 4%), and 37 non-synonymous mutations in CHIKV E2 (total mutation rate 30/425 = 7%). It is worth noting that the mutation of CHIKV E2 has changed more than that of CHIKV E1 between 2014 and 2019. CHIKV E2 screened out common mutation sites, and the results showed that there are five common mutation sites, namely S119G, L182M, G206D, S300N, and A345T. Eighty-three percent of the CHIKV E2 receptor binding domain mutations in the American strains, namely T3I and N6H, may cause immune escape.

We constructed a new luciferase immunosorbent assay using CHIKV E2 antigen and mutant E2 antigen to test the serum of the CHIKV infected patients, and the detected samples and dilution ratios were different. The T3I, N6H, S119G, L182M, G206D, S300N, and A345T mutations in CHIKV E2 could explain these differences between patients.

**IMPORTANCE:** CHIKV originated in Africa more than 500 years ago, with a common lineage dividing into two distinct branches called West Africa (West African, WA) and East/Central/South Africa (East/Central/South African, ECSA). Moreover, the E1-A226V mutation enhanced the replication and transmission capacity of CHIKV in *Aedes albopictus*. However, it should be noted that E1-A226V does not explain the CHIKV epidemic that has occurred in the Americas in recent years. We analyzed CHIKV samples in the 2014-2019, and the results showed that the mutation of CHIKV E1 protein occurred was not mainly concentrated in E1-A226V, but concentrated in M343I. This also means that CHIKV constantly mutates in the natural environment and is formed under natural conditions. Analysis of the common mutation sites of CHIKV E2 showed that there were seven common mutation sites S119G/L182M/ G206D /S300N/A345T, and the new mutation of CHIKV E2 may affect serological antibody detection.

## INTRODUCTION

Chikungunya is a single-stranded RNA virus, a member of the *Alphavirus* genus of the family *Togaviridae*, which is transmitted by mosquitoes. No longer confined to tropical developing countries, CHIKV has recently erupted in various geographical areas around the globe, including the United States and Europe. CHIKV infection has re-emerged as a significant infectious disease, causing a usually self-limiting illness characterized by sudden onset of high fever, severe joint pain and rash. CHIKV disease is usually associated with chronic and disabling arthritis. As a result, the massive CHIKV epidemic has had a severe impact on the economy, highlighting the significant public health threat posed by CHIKV infection (1).

Each CHIKV spherical virus particle is about 70 nm in diameter and consists of an 11.8 kbp positive-sense RNA genome, surrounded by a capsid (C) protein and a host cell-derived lipid bilayer, which is mixed with heterodimers of envelope proteins E1 and E2. CHIKV genome size is 12 kb, divided into two coding regions ORF (Open Reading Frame, ORF), each containing an ORF encoding nine proteins. They were divided into five structural proteins (C, E3, E2, 6K and E1) and four non-structural proteins (nsP1-4) (2).

Envelope proteins E2 and E1 play important roles in virus binding to host cell membrane and subsequent cell invasion, respectively. E1 and E2 glycoproteins form heterodimers on the surface, where E2 is primarily located above E1 and is thought to interact with cell receptors. The E1 glycoprotein promotes endosomal fusion and capsid release into the cytoplasm after CHIKV transport to the early endosomes. E2 envelope glycoprotein is an important binding site for neutralizing antibodies and mediates the binding of E1 to receptors and adhesion factors on the cell membrane (3,4,5).

After human infection with CHIKV, IgM levels are detected within 5-7 days of symptom onset, peak a few weeks after infection, and begin to diminish in the following months. IgG response can be detected about 7-10 days after onset, usually after the viremia has been cleared. Human IgM and IgG antibodies broadly and effectively neutralize CHIKV, with the most effective neutralizing antibodies targeting domains A and B of the E2 glycoprotein, and antibodies targeting domains B generally exhibit extensive neutralization against CHIKV and other related α virus strains. Rather than individual E1 or E2, IgM preferentially binds and targets epitopes on e1-E2 glycoproteins (6). This provides evidence of early antibody response to CHIKV and has implications for the design of diagnostic serology detection.

The distal viral membranes of domains A and B are the main surface exposure epitopes recognized by IgG antibodies. Most CHIKV IgG antigens are conformational epitopes in E2 space that are linear or discontinuous in conformation (7-9).Highly exposed epitopes in the extracellular domain of CHIKV E2 have important antigenicity for mAb binding and neutralization (10-13).Whether in vitro or in vivo, antibodies targeting the E2 neutralization site can effectively combat CHIKV infection, and this region may be an important area for binding to CHIKV antibodies.

The emergence of spontaneous mutations in the CHIKV genome has resulted in enhanced geographic spread across the globe, which may lead to CHIKV adaptation to different vector species. A global detection system capable of detecting the dynamic variation and transmissibility of the virus is critical. This can only be achieved by combining an efficient public health system with sensitive and specific diagnostic tools that global laboratories can use to rapidly identify infected patients and virus variants.

## RESULTS

### Phylogenetic analysis

Phylogenetic analysis using the maximum-likelihood method revealed that CHIKV is still clustered according to geographic location, and every epidemic may be related to adaptive mutations produced by local mosquitoes. Bootstrap values (>90) are indicated at major nodes. The CHIKV that occurs in the Pacific Islands and the Americas is different from the CHIKV that occurs in Africa. The Pacific Islands, the Americas and Asian strain have more variation (FIG 1). The virus strains currently occurring in all regions exhibit local aggregation.

**FIG 1.**
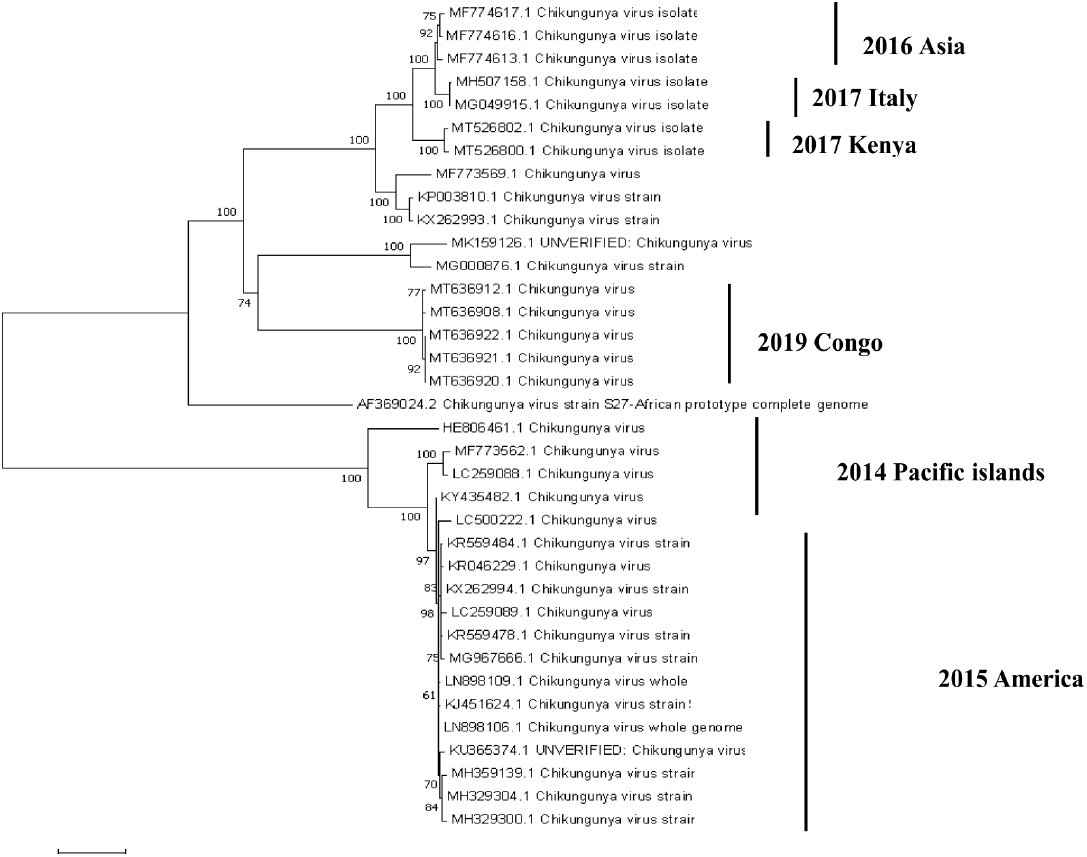
Maximum likelihood tree based on the nucleotide sequence of 31 CHIKV genes that occurred from 2014 to 2019. rap values (>90) are indicated at major nodes. Each strain is represented by its GenBank accession number, year of ion, and country of origin.

Compared with the reference CHIKV sequence (NC_004162), CHIKV E1 19 non-synonymous mutations were found in 31 sequences (the total mutation rate was 20/499 = 4%), and we found 37 non-synonymous mutations in CHIKV E2 (total mutation rate is 30/425 = 7%). From 2014 to 2019, CHIKV E2 mutations have changed more than E1 mutations.

### CHIKV E1 mutation site analysis

We analyzed the situation of CHIKV E1 non-synonymous mutations in the 2014 Pacific, 2015 Americas, 2017 Italy, 2017 Kenya, and 2019 Congo region leading to mutations in amino acid positions with mutation rates of 1.4%, 2.8%, 2.4%, 1.6%, and 1%, respectively (Table 3).

Using a Venn diagram, CHIKV E1 was screened out for common mutation sites, and one common mutation site was M343I (FIG 2). Most mutation sites belong to the American strains, and the mutation rate is 2.8%. We focused our attention on the described adaptive mutation of E1-A226V of CHIKV, but A226V was not obvious in our study. The 2015 American strain has the highest mutation rate and is currently the most widely distributed. Today, the American strain is still popular in some American countries, such as the United States. The most frequent mutations were L26M, T51M, A53T, M58L, N139S, A165T, T212A, K278E, M343I, and P371S, and more than 50% of the mutations were concentrated in the 2014 Pacific strain(FIG 3).

**FIG 2.**
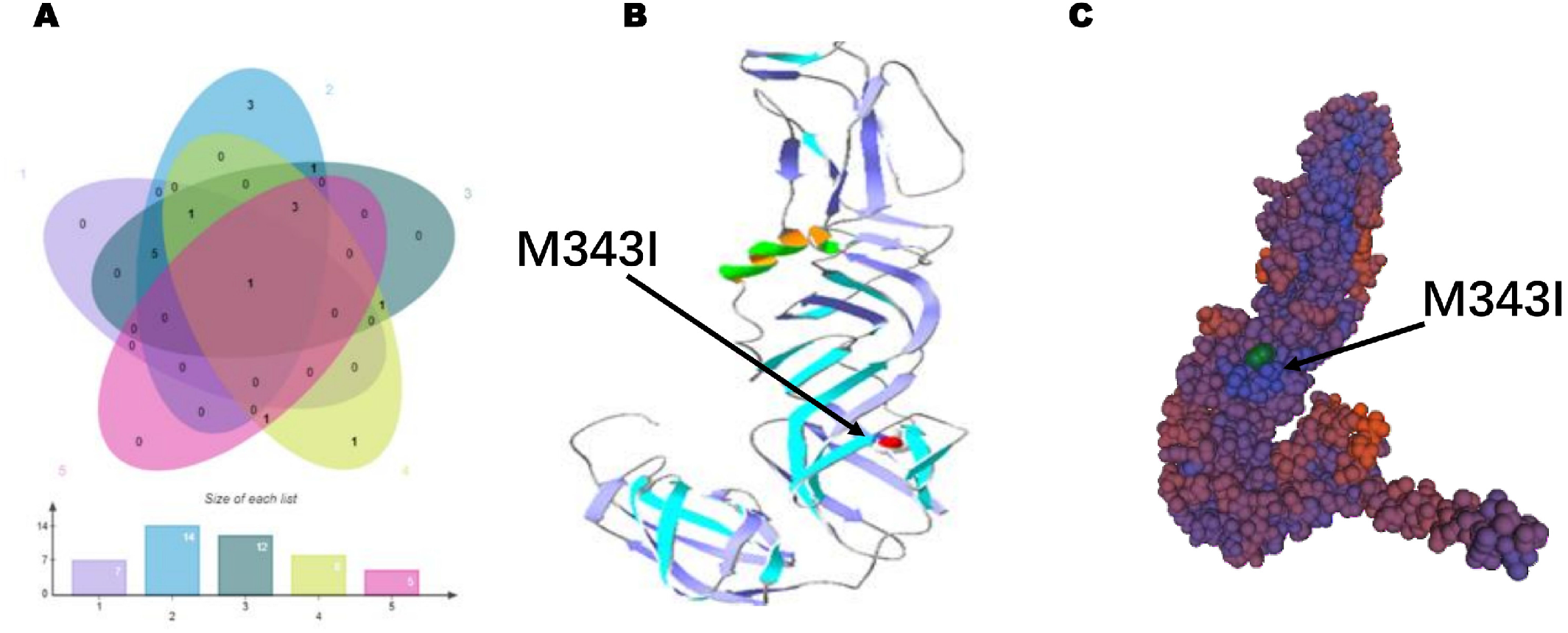
**A** Venn diagram analysis showed that CHIKV El had a common mutation site M343I. **B** CHIKV El M343I space folding structure diagram; **C** The green site is the mutation site, and the rotated site CHIKV El M343I and its protein structure model

**FIG 3.**
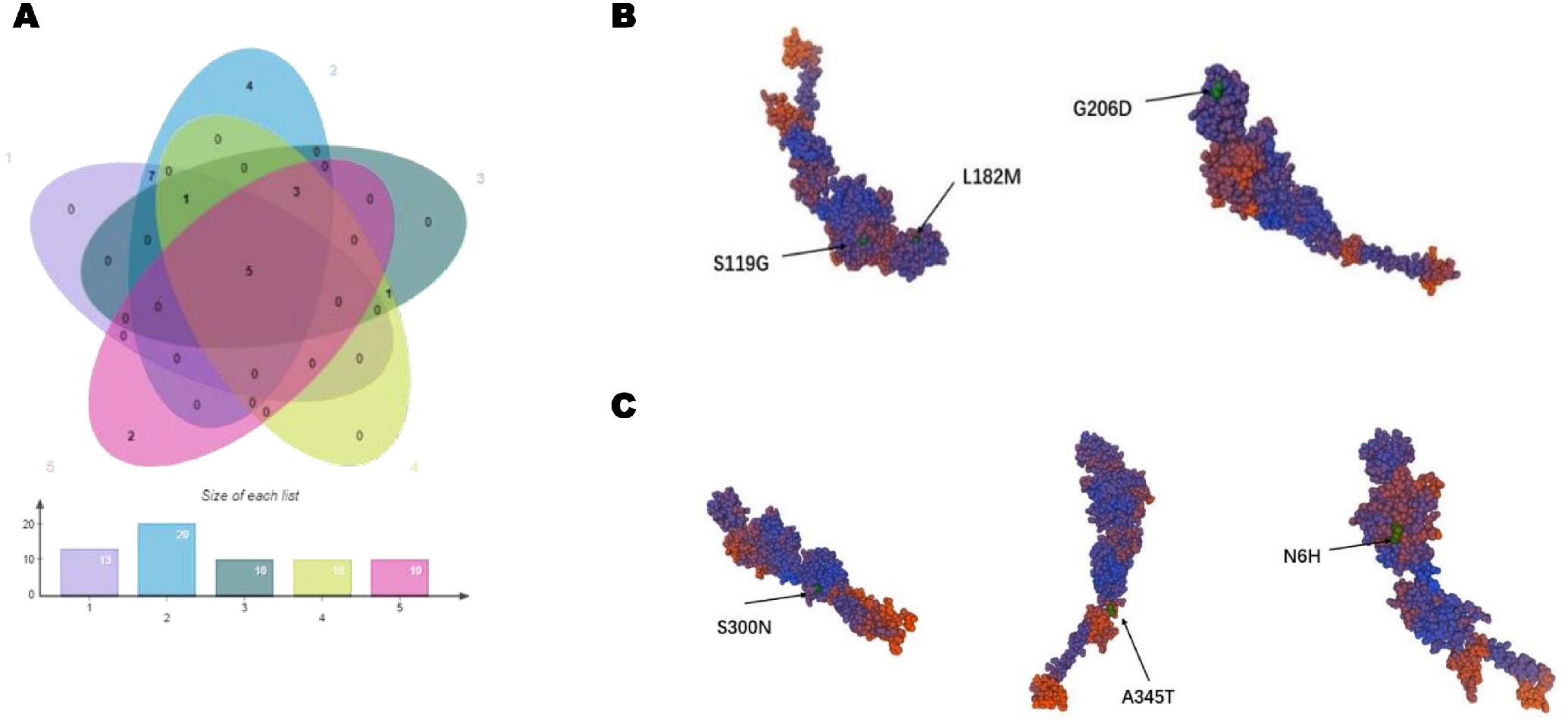
**A** venn diagram analysis showed that CHIKV E2 had 5 common mutation sites NH6, SI19G, Ll82M, G206D, S300N, A345T. **B** The green site of CHIKV E2 is the mutation site, and the rotated site CHIKV-E2 NH6/Sll9G/Ll82M/G206D/S300N/A345T and its protein structure model.

The non-synonymous mutation rates of CHIKV E1 in the 2014 Pacific, 2015 Americas, 2017 Italy, 2017 Kenya, and 2019 Congo region were 2.9%, 3.3%, 2.65%, 2.1%, and 1.8%, respectively (Table 4).

### CHIKV E2 mutation site analysis

Using the Venn diagram, CHIKV E2 was screened out for common mutation sites, and five common mutation sites were found, namely S119G, L182M, G206D, S300N, and A345T. The most mutated strains belong to the American strains, with a mutation rate of 3.3%. The American strains are still circulating in the United States today.

The antibody response to the CHIKV capsid protein, E2, and E3 glycoproteins, and non-structural protein (nsP3) protein determinants was high but gradually diminished. Only the reaction to the glycoprotein E2 could still be detected after 21 months. The sustained response to the N-terminal region of E2 glycoprotein (between amino acids 2800 and 2818) is important for sero-epidemiology studies (14). High levels of antibodies against the epitope E2EP3 detected during the acute phase of CHIKV proved to be associated with long-term clinical protection. Antibodies significantly recognize the major linear epitopes of E2 glycoproteins (E2 P1-1 and P1-2), which correspond to linear E2EP3 epitopes previously identified from patient plasma. In plasma samples obtained early in the recovery period after CHIKV infection, the naturally occurring IgG response was predominantly based on IgG3 antibodies to a single linear epitope “E2EP3” (15-17).

E2EP3 is located at the N-terminus of E2 glycoprotein and is mainly exposed to the viral envelope. The “E2EP3” specific antibody has a neutralizing effect, and when removed from the plasma the titer of CHIKV-specific antibody can be reduced by up to 80%. These results suggest that the Arch1 region is a key region for detecting CHIKV anti-IgG antibodies, and its mutation may affect antibody detection.

The CHIKV receptor binding domain (RBD) is an important domain that binds to antibodies. Amino acid mutations located in the RBD domain of CHIKV E2 glycoprotein will cause E2 mutations to a greater extent, resulting in decreased antibody detection capabilities and even immune escape. The T3I and N6H mutations in the RBD domain of CHIKV E2 occurred in the 2014 Pacific/2015 Americas, and 83% of the American strains also had the T3I and N6H mutations.

After data analysis, we believe that T3I, N6H, S119G, L182M, G206D, S300N, and A345T sites are the key sites (FIG 4).

**FIG 4.**
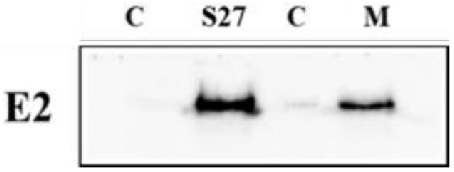
C:control group. S27: Africa Chikungunya S27 strain. M: Chikungunya virus 7-site mutant.

### CHIKV E2 sequence key-site induced mutations

We studied the mutations at the sites mentioned above. Adaptive CHIKV E1 and E2 mutations lead to the decline of antibody detection ability, which is formed under natural conditions and requires further attention.

The base sequence of Africa Chikungunya S27 strain E2 and the base sequence corresponding to seven amino acid site mutations (T3I, N6H, S119G, L182M, G206D, S300N, and A345T) were inserted into the pNLF1-N plasmid of Nano luciferase, and the plasmid was transfected into cells for protein expression. WB was used to detect proteins of CHIKV S27 and mutant strains (FIG 5). The nano-antigen labeled with fluorescein was combined with the antibody to observe IgG resistance’s dilution ratio and detection ability. S119G, L182M, G206D, S300N, and A345T are located in domain A, domain D, domain D, domain C, and E2 stem, respectively (FIG 6).

**FIG 5.**
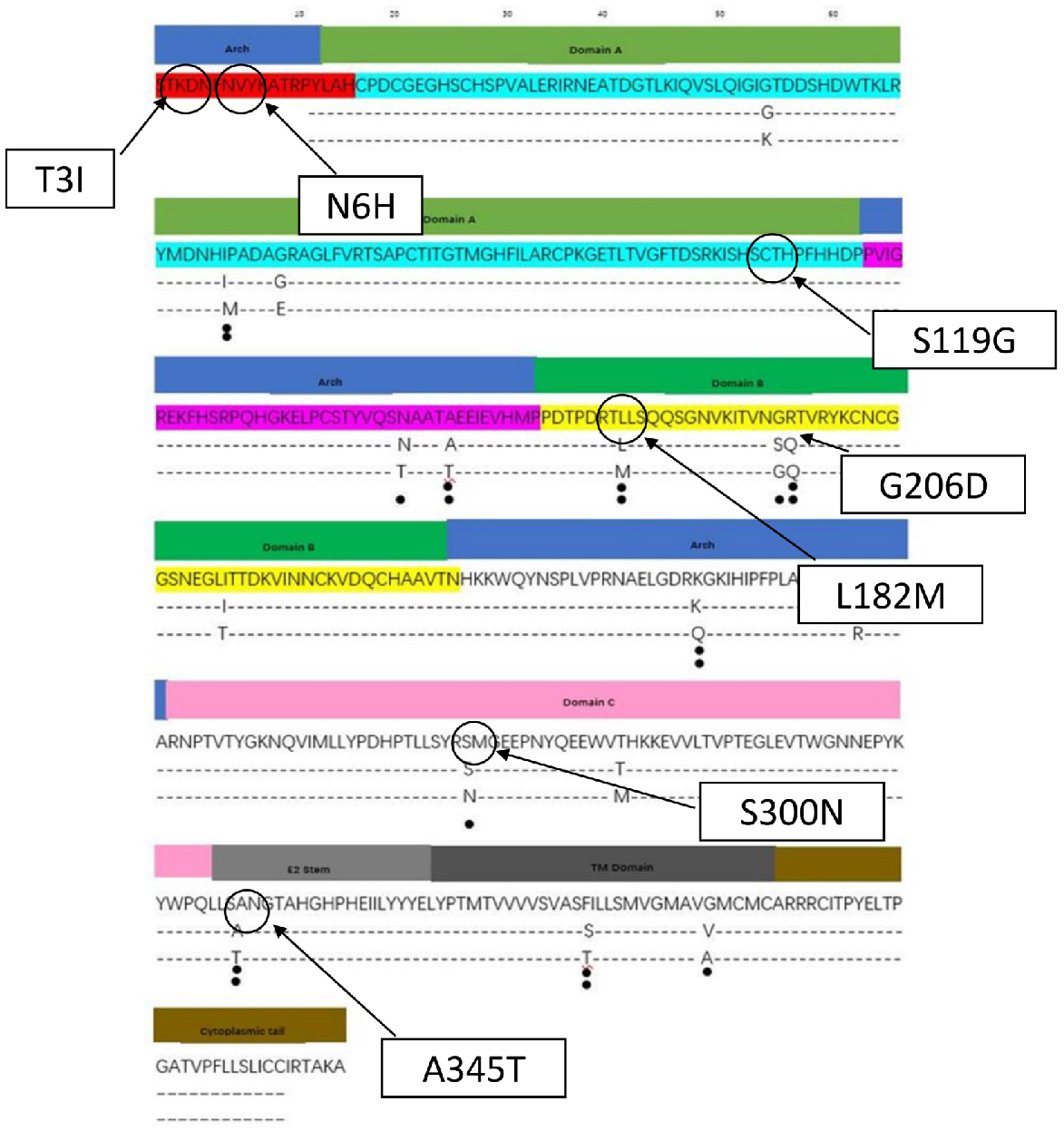
Seven mutation sites: The T31/N6H in Archl2; Sll9G resides in Domain A3; L182M /G206D resides in Domain D4; The S3OON resides in Domain C5; A345T is located in E2 stem

**FIG 6.**
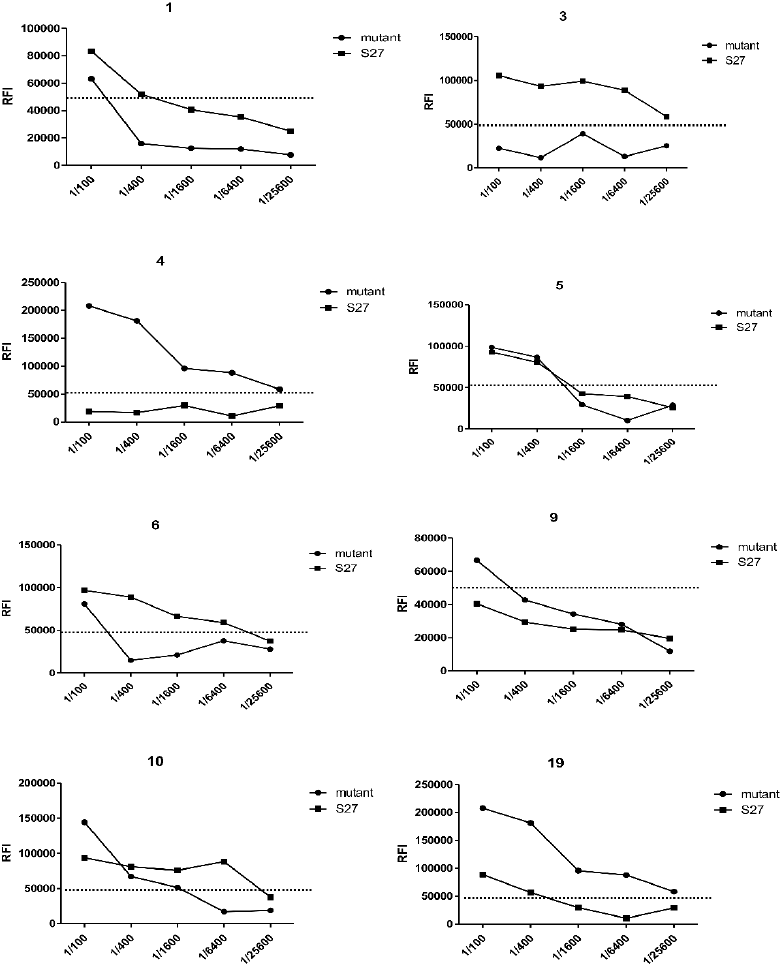
CHlKV S27 E2 antigen can detect positive serum CHIKV IgG antibody numbered 1,3,5,6, 10, 19; CHIKV E2 mutant showed positive CHIKV IgG antibody and the numbers were 1,4,5,6,9,l0,19; ln sample 6, the detection dilution ratio of CHIKV E2 mutant was 1:200, and the detection dilution ratio of CHlKV S27 E2 antigen was 1:6,400; ln sample 10, the detection dilution ratio ofCHIKV S27 E2 antigen was J:1600, and that of CHIKY E2 mutant was 1:25600; ln sample 19, the detection dilution ratio of CHIKV S27 E2 antigen was 1:1600, and that of CHIKV E2 mutant was **J** :25600

### CHIKV anti-IgG antibody was detected by fluorescein immunosorbent assay

The control samples’ nucleic acid, serum IgM, and IgG antibodies were all negative (commercial kits). Panel A (reference nos. 21–42) included serum samples from 21 healthy controls with no past infection of CHIKV as negative controls, confirmed by the absence of antibodies by a serum neutralization assay.

Twenty samples were obtained from patients diagnosed with CHIKV infection (reference nos. 21–42). Samples were collected from patient sera collected at the Guangdong Provincial Center for Disease Control and Prevention during the 2010 CHIKV outbreak; acute samples were collected from viremic patients between days 2 and 11 after disease onset and eight months later. Convalescent serum was collected. In a previous study, real-time PCR targeting the E1 region was used to quantify viral load and, in conjunction with clinical symptoms, confirm CHIKV infection (Table 2).

LISA detected the RFI of serum of normal patients. We comprehensively analyzed the normal subjects’ RFI and CHIKV infected people. To ensure specificity, we compared the mean (SEM) + standard error in the CHIKV infected population, approximately twice the control group’s RFI value. We defined the RFI cut-off value of positive anti-CHIKV IgG antibody as 5 × 10^4^.

Antigens of African CHIKV S27 virus strain and CHIKV mutant E2 were used to detect the serum of CHIKV infected patients. The experimental results showed that the CHIKV S27 strain could detect positive serum CHIKV anti-IgG antibody (samples 1, 3, 5, 6, 10, and 19), while the CHIKV mutant strain could not detect sample 3. Joint-detection was found for samples 1, 6, 10, and 19. The dilution ratio of antigens detected by CHIKV S27 and CHIKV E2 mutant was different, and samples 6, 10, and 19 displayed the greatest difference. In sample 6, the detection dilution ratio of CHIKV E2 mutant was 1:200, and the detection dilution ratio of CHIKV S27 E2 antigen was 1:6,400. In sample 10, the detection dilution ratio of CHIKV S27 E2 antigen was 1:1,600, and that of CHIKV E2 mutant was 1:25,600. In sample 19, the detection dilution ratio of CHIKV S27 E2 antigen was 1:1,600, and that of CHIKV E2 mutant was 1:25,600.

The detection dilution ratio of the mutant strain was lower than that of the CHIKV S27 strain (nos. 4 and 9). The serum anti-CHIKV IgG antibody could be detected in the CHIKV mutant strain, while the serum anti-CHIKV IgG antibody could not be detected in the CHIKV S27 strain (FIG 6). The above sites were mutated, and the CHIKV E2 mutation led to decreased antibody detection ability, both of which were formed under natural conditions, requiring further attention.

## DISCUSSION

The continuous mutation of the CHIKV is closely related to its increasing global spread. Currently, the molecular mechanisms by which the mutated CHIKV interacts with host cells are not fully explained. However, continuous mutations in CHIKV improve its adaptability to *Aedes aegypti* or *Aedes albopictus*, which leads to an increase in the ability of the virus to spread among people and to expand the area where the virus can spread (18). The emergence of ECSA-E1-A226V may be one of the reasons for the recent expansion of the epidemic in Italy and China (19). Mutations in the surface envelope glycoproteins E1 at position 226 (E1-A226V) have been shown to enhance CHIKV replication and transmission in *Aedes mosquitoes* (20,21). In addition, it is important to note that none of the characteristics of ECSA-E1-A226V can explain the scale of the recent outbreak in the Americas, as the E1-A226V mutation was not found in samples collected during the Americas outbreak (22). Our results also validate Feng’s results that the mutation of the CHIKV E1 protein occurring during 2014–2019 was not mainly concentrated in E1-A226V. We collected 20 samples (reference nos. 1–20) from patients who visited the Guangdong Provincial Center for Disease Control and Prevention during the 2010 CHIKV outbreak. Viral loads were quantified using real-time PCR targeting the E1 region in a previous study. In the 2007 Italy outbreak, *Aedes albopictus* with ECSA-E1-A226V could develop a shortened extrinsic incubation period and higher transmission potentials (23,24). The emergence of ECSA-E1-A226V may have led to outbreaks not only in Italy but also in China (18). However, our mutation analysis in the most recent past five years showed that the E1-A226V mutation was not significant because the most common mutation site in CHIKV E1 was M343I.

In addition, it should be noted that ECSA-E1-A226V cannot explain the size of the recent outbreak in the Americas, as no mutation was found in samples collected during the outbreak in the Americas (25). Our results also validated Feng’s findings that the mutations of CHIKV E1 protein during 2014–2019 were not mainly concentrated in E1-A226V (26). When the CHIKV S27 and CHIKV mutant strain E2 antigens were detected in serum samples, there were differences in detection and dilution ratios. The possible reason for these differences is the seven mutations of CHIKV E2 antigen, namely T3I, N6H, S119G, L182M, G206D, S300N, and A345T.

Domain A is known to contain the putative RBD. Among the antibodies to the E2 epitope, most neutralizing antibodies preferentially recognize sites in region A and Arch1 and 2, while few groups recognize sites in domain B (27). These data suggest that the surface-exposed regions of domain A and Arch cause neutralizing antibody responses in humans (28,29). Highly conserved regions in domain A and Arch2 may elicit a broadly protective immune response. The T3I, N6H, and S119G sites may play a more important role in antibody detection than other sites.

## MATERIALS AND METHODS

### Sample Information

As shown in the figure, we analyzed the entire Chikungunya gene sequence in the Pacific Islands, Americas, Asia, Europe/Africa, and Africa from 2014 to 2019 and selected 31 CHIKV sequences (Table 1). In this study, 31 CHIKV sequence numbers were analyzed: AF 369024, MF773562, LC259088, MF773569, KU365374, MK159126, LC500222, KY435482, KR559478, LN898109, MH329304, MG000876, LC259089, KJ451624, KX262994, KR559484, KR046229, LN898106, MH329300, MH359139, MG967666, MH507158, MG049915, KP003810, KX262993, MT636922, MT636921, MT636920, MT636912, and MT636908. The original strain is the Africa Chikungunya S27 strain.

**Table 1.** Chikungunya fever occurred in major areas in 2014-2019

| **Year** | **Country** |
| --- | --- |
| 2014 Pacific Islands | 1.Papua New Guinea 2. Kiribati 3. Mauritius 4. Samoa<br>5. Tonga |
| 2015 Americas | 1.Brazil 2. Cuba 3. Dominica 4. Ecuador 5. El Salvador<br>6. French Guiana 7. Grenada 8. Guadeloupe 9. Haiti10.<br>Martinique 11. Montserrat 12. Puerto Rico13. Saint<br>Kitts and Nevis 14. Saint Lucia 15. Suriname 16. Tri-<br>nidad and Tobago 17. Turks and Caicos Islands Ven-<br>ezuela 19. Aruba 20. Anguilla 21. Antigua and Barbuda<br>Colombia 23. Cayman Islands 24. British Virgin Islands |
| 2016 Asia | 1.Pakistan |
| 2017 Europe/Africa | 1. Italy 2. Kenya |
| 2019 Africa | 1.Congo 2. People's Republic of Congo |

**Table 2.** Detection sample information.

| Group | Reference<br>no | Source (human) |  | CHIKV-RT-PCR-E1 | ELISA(IgG) | LISA(IgG) |
| --- | --- | --- | --- | --- | --- | --- |
|  |  | Region | Notes |  |  |  |
| CHIKV-serum<br>(n=20) | 1-2 | China |  | + | - | + |
|  | 3 | China | about 8 months | + | - | + |
|  | 4-7 | China |  | + | - | + |
|  | 8 | China | 3 d, acute phase | + | - | + |
|  | 9 | China | 3 d, acute phase | + | - | + |
|  | 10 | China | 11 d, acute | + | - | + |
|  | 11 | China | 2 d, acute phase | + | - | + |
|  | 12 | China |  | + | - | + |
|  | 13 | China | 2 d, acute phase | + | - | + |
|  | 14 | China | 2 d, acute phase | + | - | + |
|  | 15 | China | 6 d, acute phase | + | - | + |
|  | 16 | China | 3 d, acute phase | + | + | + |
|  | 17 | China |  | + | + | + |
|  | 18-20 | China |  | + | - | + |
| Control<br>(n=20) | 21-42 | China |  | - | - | - |

**TABLE 3.** Comparison of amino acid substitutions between the 31 partial E1 sequences in the present study with the Chikungunya S27 (NC_004162)

| GenBank | Amino acid position in Chikungunya S27 strain (NC_004162) |  |  |  |  |  |  |  |  |  |  |  |  |  |  |  |  |  |  |  |  |  |  |  |  |  |  |  |  |
| --- | --- | --- | --- | --- | --- | --- | --- | --- | --- | --- | --- | --- | --- | --- | --- | --- | --- | --- | --- | --- | --- | --- | --- | --- | --- | --- | --- | --- | --- |
| accession number | 14 | 26 | 51 | 53 | 58 | 60 | 66 | 76 | 104 | 139 | 165 | 212 | 278 | 292 | 293 | 317 | 336 | 343 | 351 | 355 | 358 | 371 | 384 | 389 | 391 | 444 | 474 | total | mutation rate |
| NC_004162 | V | L | T | A | M | V | N | S | I | N | A | T | K | A | V | P | V | M | D | T | V | P | I | A | R | A | M |  |  |
| MF773562 | V | L | M | T | L | I | S | N | T | S | T | A | E | S | A | S | M | I | D | T | V | S | I | V | K | A | M | 7 | 0.014 |
| LC259088 | V | L | M | T | L | I | S | N | T | S | T | A | E | S | A | S | M | I | D | T | V | S | I | V | K | A | M | 7 | 0.014 |
| MF773569 | I | L | T | A | M | I | S | N | T | N | A | T | K | A | V | S | V | I | E | T | V | P | I | V | K | A | M | 6 | 0.012 |
| KU365374 | V | M | M | T | L | I | S | N | T | S | T | A | E | S | A | S | M | I | E | T | V | S | I | V | K | A | M | 10 | 0.020 |
| MK159126 | V | L | T | A | M | I | S | N | T | N | A | T | K | A | A | S | V | I | E | I | V | P | I | V | K | V | L | 4 | 0.008 |
| LC500222 | V | M | M | T | L | I | S | N | T | S | T | A | E | S | A | S | M | I | E | T | V | S | I | V | K | A | M | 9 | 0.018 |
| KY435482 | V | M | M | T | L | I | S | N | T | S | T | A | E | S | A | S | M | I | E | T | A | S | I | V | K | A | M | 10 | 0.020 |
| KR559478 | V | M | M | T | L | I | S | N | T | S | T | A | E | S | A | S | M | I | E | T | V | S | I | V | K | A | M | 9 | 0.018 |
| LN898109 | V | M | M | T | L | I | S | N | T | S | T | A | E | S | A | S | M | I | E | T | V | S | I | V | K | A | M | 9 | 0.018 |
| MH329304 | V | M | M | T | L | I | S | N | T | S | T | A | E | S | A | S | M | I | E | T | V | S | I | V | K | A | M | 8 | 0.016 |
| MG000876 | V | M | T | A | M | I | S | N | T | N | A | T | K | A | A | S | V | I | D | I | V | P | I | V | K | V | L | 6 | 0.012 |
| LC259089 | V | L | M | T | L | I | S | N | T | S | T | A | E | S | A | S | M | I | D | T | V | S | I | V | K | A | M | 7 | 0.014 |
| KJ451624 | V | M | M | T | L | I | S | N | T | S | T | A | E | S | A | S | M | I | D | T | V | S | I | V | K | A | M | 7 | 0.014 |
| KX262994 | V | M | M | T | L | I | S | N | T | S | T | A | E | S | A | S | M | I | D | T | V | S | I | V | K | A | M | 7 | 0.014 |
| KR559484 | V | M | M | T | L | I | S | N | T | S | T | A | E | S | A | S | M | I | D | T | V | S | I | V | K | A | M | 7 | 0.014 |
| KR046229 | V | M | M | T | L | I | S | N | T | S | T | A | E | S | A | S | M | I | D | T | V | S | I | V | K | A | M | 7 | 0.014 |
| LN898106 | V | M | M | T | L | I | S | N | T | S | T | A | E | S | A | S | M | I | D | T | V | S | I | V | K | A | M | 7 | 0.014 |
| MH329300 | V | M | M | T | L | I | S | N | T | S | T | A | E | S | A | S | M | I | D | T | V | S | I | V | K | A | M | 7 | 0.014 |
| MH359139 | V | M | M | T | L | I | S | N | T | S | T | A | E | S | A | S | M | I | D | T | V | S | I | V | K | A | M | 7 | 0.014 |
| MG967666 | V | M | M | T | L | I | S | N | T | S | T | A | E | S | A | S | M | I | D | T | V | S | I | V | K | A | M | 7 | 0.014 |
| MH507158 | I | M | T | A | M | I | S | N | T | N | A | T | E | A | A | S | V | I | E | T | V | P | I | V | K | A | M | 7 | 0.014 |
| MG049915 | I | L | T | A | M | I | S | N | T | N | A | T | E | A | A | S | V | I | E | T | V | P | V | V | K | A | M | 7 | 0.014 |
| KP003810 | I | L | T | A | M | I | S | N | T | N | A | T | K | A | V | S | V | I | E | T | V | P | I | V | K | A | M | 6 | 0.012 |
| KX262993 | I | L | T | A | M | I | S | N | T | N | A | T | K | A | V | S | V | I | E | T | V | P | I | V | K | A | M | 5 | 0.010 |
| MT636922 | V | L | T | A | M | I | S | N | T | N | A | T | K | A | V | S | V | I | D | T | V | P | I | V | K | A | M | 4 | 0.008 |
| MT636921 | V | L | T | A | M | I | S | N | T | N | A | T | K | A | V | S | V | I | D | T | V | P | I | V | K | A | M | 4 | 0.008 |
| MT636920 | V | L | T | A | M | I | S | N | T | N | A | T | K | A | V | S | V | I | D | T | V | P | I | V | K | A | M | 4 | 0.008 |
| MT636912 | V | L | T | A | M | I | S | N | T | N | A | T | K | A | V | S | V | I | D | T | V | P | I | V | K | A | M | 4 | 0.008 |
| MT636908 | V | L | T | A | M | I | S | N | T | N | A | T | K | A | V | S | V | I | D | T | V | P | I | V | K | A | M | 4 | 0.008 |

**TABLE 4.**
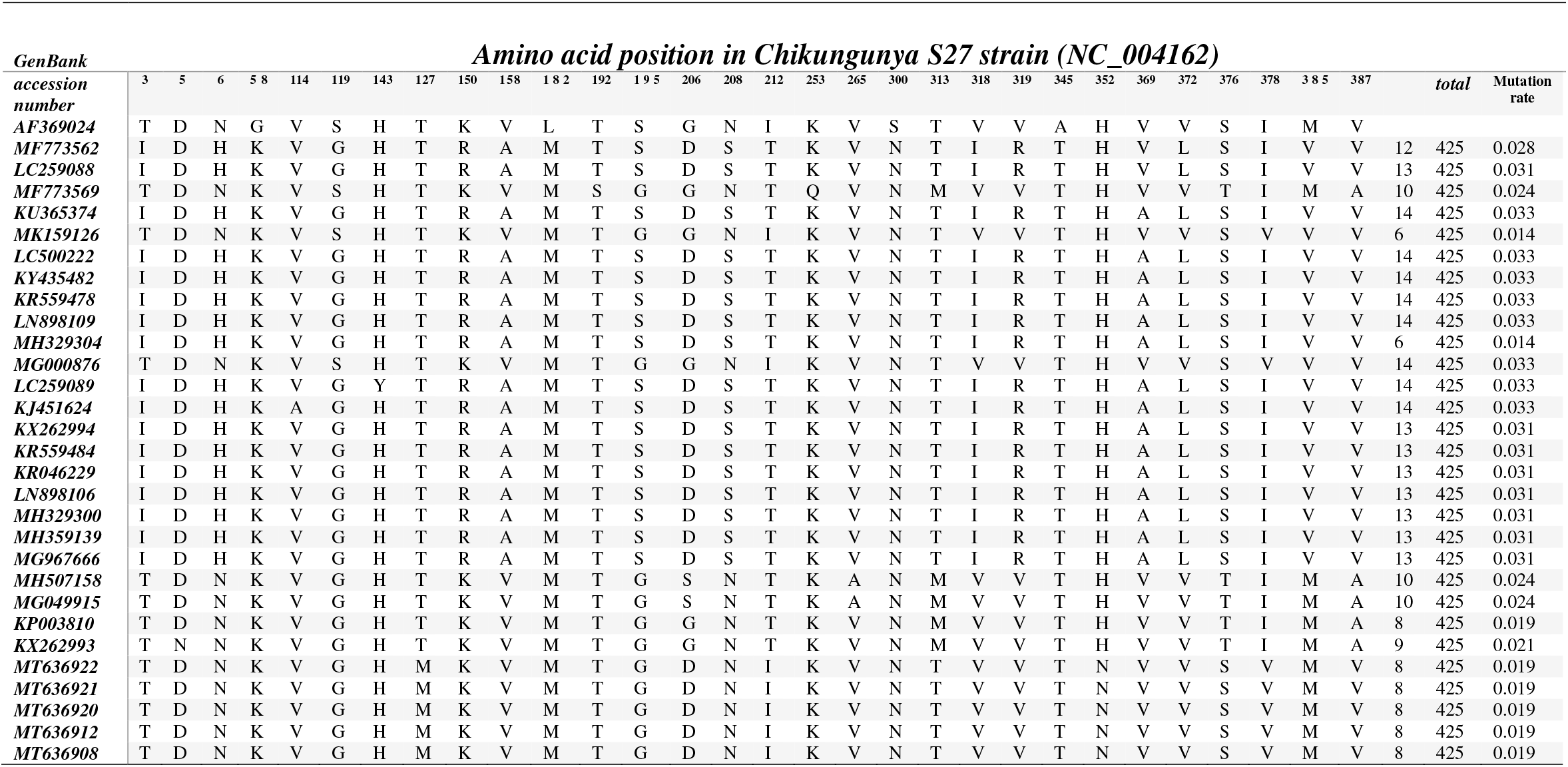
Comparison of amino acid substitutions between the 31 partial E2 sequences in the present study with the Chikungunya S27 (NC_004162

### Multiple sequence alignment

The variations at the nucleotide/amino acid level of the sequences were analyzed concerning the CHIKV prototype strain S27 (GenBank accession number: NC_0 04162). CHIKV nucleotide sequences from human specimens between 2014 and 2019 are available from NCBI through the website at https://www.ncbi.nlm.nih.gov/ and were used for further analysis. The entire CHIKV gene sequence mentioned above is the screening criteria from 2014 to 2019. Representative sequences that meet the requirements are screened out through gene comparison. All the sequences were aligned using the ClustalX algorithm in the Mega program. Sequences that did not adequately cover the region of interest were excluded.

### Assembly of Genome Sequences and Sequence Analysis

Contig assembly was performed independently by distinct operators and software, using Bioedit. Alignments of nucleotide and amino acid (aa) sequences against selected CHIKV sequences were performed with the ClustalW version. Neighbor-joining trees were constructed using MEGA version 7. The maximum likelihood tree is based on the nucleotide sequence of 31 CHIKV genes from 2014 to 2019. Bootstrap values (> 90) are indicated at major nodes. Each strain is represented by its GenBank accession number, year of collection, and country of origin.

### 3D Structure Modelling

The crystallographic structure of the ectodomain of the glycoprotein E2 of CHIKV at neutral pH was used as a template to model and analyze the amino acid mutations of the 2014–2019 isolates. The 3D structure figure was prepared using the program SWISS MODEL.

### Plasmid construction

A synthetic sequence of CHIKV E2 sequence (GenBank accession number: NC_004162) was generated (Sangon Biotech, China) and inserted into the pNLF1-N plasmid with luciferase to construct a recombinant plasmid. The recombinant plasmid was transfected into the human embryonic kidney (HEK) 293T cells, and the protein expression was detected by western blot (WB). Similarly, the mutations were inserted into the pNLF1-N plasmid. The mutation site is CHIKV E2 T3I/N6H/S119G/L182M/G206D/S300N/A345T.

### Cell culture and transfection

Cryopreservation tubes containing HEK 293T cells were removed from liquid nitrogen and immersed directly in a water bath at a constant temperature of 37°C, without shaking, to promote the rapid thawing of the cells. The 293T cells were resuscitated in DMEM (dulbecco’s modified eagle medium, DMEM) medium containing 10% fetal bovine serum (Gibco, New Zealand), and the cell culture flask was placed in an incubator at 37°C. The luciferase recombinant plasmid was transfected into 293T cells with lip 3000 transfection reagent to express the luciferase target antigen. After the reaction system was prepared, it was still for 10–15 minutes. After changing the medium, the reaction medium was transferred into 293 T cells, cultured in a CO_2_ incubator at 37°C for 48 h, and collected.

### Western Blot

The cell lysate proteins were resolved with 10% SDS(Sodium dodecyl sulfate, SDS)-PAGE under non-reducing or reducing conditions and electro-transferred onto a nitrocellulose (NC) membrane (China). The membrane was blocked with 5% skim milk in 0.05% phosphate-buffered saline (PBS)-Tween 20 (PBST). For His-Tag detection, the protein expression recombinant CHIKV proteins and viral antigens were evaluated at 1:5,000 and 1:10,000 dilutions. The HRP (Horseradish Peroxidase, HRP) ECL (Electro-Chemi-Luminescence, ECL) luminescence method was used, and the color-developing solution was mixed with solution A and solution B at a 1:1 ratio before adding it to the NC film. After placing the film into the black box, the film was colored and stored.

### CHIKV immune serum panels

This study used two panels of serum samples. Panel A (reference nos. 21–42) included serum samples from 21 healthy controls with no past infection of CHIKV as negative controls, confirmed by the absence of antibodies by serum neutralization assay. Panel B comprised 20 (reference nos. 1–20) samples collected from patients attending Guangdong Center for Disease Control and Prevention, China, during the 2010 outbreak of CHIKV. These were acute samples collected from viremic patients between day two and day 11 after disease onset, and convalescent sera collected eight months later. The viral loads were quantified in a previous study using real-time PCR(Polymerase Chain Reaction, PCR) targeting the E1 region. The anti-CHIKV IgG antibody in the serum was detected by the luciferase immunosorbent assay (LISA) detection method, and they were all positive.

### Development of LISA based on different E-luciferase fusion proteins

Protein G (5 μg/ml, 50 μl/well; Genscript, China, Z02007) was diluted to 5 μg/ml with PBS (0.01 M, pH 7.4), added to the whiteboard of 96-well flat-bottomed luminometer plates (Corning Costar, Shanghai China 3922) at 100 μL per well, and coated at 4°C overnight. First, 5% skim milk in PBS was used as a sealing solution. The coated board was then washed with PBST (0.05% Tween PBS) after 1-h incubation of the sealing solution. Next, 300 μL of sealing solution was added to each well, and the wells were incubated at 37°C in an incubator for 1 h. Following incubation, the wells were washed with PBST three times and dried as much as possible after the final wash. Next, the sample was diluted with 2% skim milk (Sangon Biotech, Shanghai China, A600669-0250) and added to the whiteboard at 100 μL per well. Positive, negative, and blank controls were set in each experiment. Next, the whiteboard was incubated at 37°C for 1 h. Following incubation, the wells were washed with PBST five times and dried as much as possible after the final wash. The supernatant of luciferase target antigen was then diluted with 2% skim milk (CHIKV-E antigen diluted 1,000-fold). Next, 50 μL was added to each well and incubated in a 37°C incubator for 1 h. Following incubation, the wells were washed with PBST five times and dried as much as possible after the final wash. Finally, according to the instructions, 50 μL of luciferase substrate (Promega, Shanghai China, N1120) was added to each well. The fluorescence value was measured within 2 h using a fluorophotometer. All steps were performed according to the manufacturer’s instructions.

To avoid differences in transfection efficiency and protein expression of different batches of preparations, we measured the luciferase activity of crude cell lysates to determine the relative fluorescence intensity (RFI), which is usually between 10^4^ and 10^6^. Therefore, the CHIKV antigen was always added in each reaction with 10^5^ RFI for LISA. In addition, we included positive and negative controls in each reaction plate to keep the results consistent and reproducible.

### Statistical analysis

Data are presented as means ± standard deviation (SD) or means ± standard error of the mean (SEM). As stated in the figure labels, differences between groups and controls were analyzed using appropriate statistical tests. A *P*-value < 0.05 was considered significant. Statistical analyses were performed using GraphPad Prism 5.04 software and SPSS 20. *P*-values were determined by unpaired two-tailed t-test, with \**P* < 0.05 and \*\**P* < 0.01.

## Acknowledgments

This work was supported by the Natural Science Foundation of Guangdong Province (number 2018B030311063) and Guangdong Science and Technology Program Key Projects (number 2021B1212030014).

We thank LetPub (www.letpub.com) for its linguistic assistance during the preparation of this manuscript.

We declare no conflicts of interest.

